# R2ROC: An efficient method of comparing two or more correlated AUC from out-of-sample prediction using polygenic scores

**DOI:** 10.1101/2023.08.01.551571

**Authors:** Md. Moksedul Momin, Naomi R Wray, S. Hong Lee

## Abstract

Polygenic risk scores (PRSs) enable early prediction of disease risk. Evaluating PRS performance for binary traits commonly relies on the area under the receiver operating characteristic curve (AUC). However, the widely used DeLong’s method for comparative significance tests suffer from limitations, including computational time and the lack of a one-to-one mapping between test statistics based on AUC and *R*^2^. To overcome these limitations, we propose a novel approach that leverages the Delta method to derive the variance and covariance of AUC values, enabling a comprehensive and efficient comparative significance test. Our approach offers notable advantages over DeLong’s method, including reduced computation time (up to 150-fold), making it suitable for large-scale analyses and ideal for integration into machine learning frameworks. Furthermore, our method allows for a direct one-to-one mapping between AUC and *R*^2^ values for comparative significance tests, providing enhanced insights into the relationship between these measures and facilitating their interpretation. We validated our proposed approach through simulations and applied it to real data comparing PRSs for diabetes and coronary artery disease (CAD) prediction in a cohort of 28,880 European individuals. The PRSs were derived using genome-wide association study summary statistics from two distinct sources. Our approach enabled a comprehensive and informative comparison of the PRSs, shedding light on their respective predictive abilities for diabetes and CAD. This advancement contributes to the assessment of genetic risk factors and personalized disease prediction, supporting better healthcare decision-making.

## Introduction

Complex diseases are affected by a multitude of risk factors, including polygenic effects^1-3^. To quantitatively evaluate the impact of these polygenic effects on future disease risk at both the individual and population levels^4, 5^, genetic profile analysis serves as a valuable tool. By analyzing genetic profiles and utilizing polygenic scores, individuals can gain meaningful insights that empower them to make informed decisions regarding their healthcare and overall well-being^6-8^. Polygenic scores have emerged as a widely employed approach in this context^9^, aiding in the assessment of genetic risk factors and personalized disease prediction.

Genome-wide association studies (GWASs) have revolutionized the estimation of PRSs for individual risk prediction using genetic data^4, 9-14^. Typically, these studies involve estimating the effects of genome-wide single-nucleotide polymorphisms (SNPs) associated with complex traits using a discovery dataset^15^, which are then applied to an independent target dataset. In the target dataset, each individual’s weighted genotypic coefficients are calculated based on the projected SNP effects, resulting in the derivation of PRSs. These PRSs are subsequently correlated with the outcome of interest, such as disease status, to assess the accuracy of risk prediction. When evaluating binary disease traits, the performance of the PRS is commonly quantified using the area under the receiver operating characteristic curve (AUC). The AUC provides a measure of the model’s ability to discriminate between affected and unaffected individuals based on the PRS.

DeLong’s test is a statistical test used to compare the AUC of two or more correlated predictors, proposed by Emily DeLong and colleagues in 1988^16^. The DeLong’s test is a non-parametric approach that does not assume any specific distribution of the data. It compares the differences between paired AUC values and their standard errors to calculate a p-value. A p-value below the chosen significance level indicates that the difference in AUC values is statistically significant. This test is widely employed in medical research and diagnostic testing to compare the performance of different tests in disease diagnosis or outcome prediction.

Despite its usefulness, DeLong’s approach has certain drawbacks. One limitation is its computational burden, which grows quadratically with the sample sizes, making it impractical for large sample sizes. Although a modified version of DeLong’s test^17^ has been proposed, which approximates the covariance using a closed-form formula based on the Spearman correlation coefficient between the predicted probabilities of the compared models (implemented in pROC R-package^18^), it still requires considerable time. This time-consuming nature hampers its application when involving large-scale or iterative analyses such as those encountered in machine learning frameworks. Furthermore, while there exists a non-linear one-to-one mapping between AUC and *R*^2^ for each metric individually, as demonstrated by Wray et al.^11^ and Lee et al.^19^, such a direct mapping does not exist when comparing the significance of AUC and *R*^2^ values in the context of comparative tests. *R*^2^ assesses the goodness of fit and the amount of explained variation in regression models, while AUC evaluates the classification performance and the ability to discriminate between classes in binary classification problems. Therefore, establishing a direct one-to-one mapping between AUC and *R*^2^ values for comparative significance tests would be desirable, as it would enhance the interpretability of the results and provide insights into the relationship between these measures. This unified approach would provide a more holistic understanding of the models’ performance and improve our ability to evaluate and interpret the results accurately.

In this study, we employ the Delta method to derive the variance and covariance of AUC values, which can be directly transformed from *R*^2^ measures. This enables the development of a comparative significance test statistic for evaluating two or more correlated AUC values. Our proposed approach offers notable advantages over the traditional DeLong’s method^16^, particularly in terms of computing time, where our method performs more than 100 times faster. Moreover, our novel approach establishes a direct relationship between AUC and *R*^2^ values in the context of comparative significance tests. This direct mapping between AUC and *R*^2^ provides a comprehensive understanding of the predictive performance captured by AUC and the explained variance captured by *R*^2^, thereby enriching the analysis and improving the interpretation of results. To demonstrate the effectiveness of our approach, we apply it to real data from a comparison of PRSs calculated in 28,880 European individuals. The PRSs are derived using GWAS summary statistics for diabetes from two distinct sources: the UK Biobank (UKBB) and Biobank Japan (BBJ). By utilizing our proposed approach, we are able to perform a comprehensive and informative comparison of these PRSs, shedding light on their respective predictive abilities for diabetes.

## Material and methods

### Area under ROC (AUC)

AUC is a widely used metric in various fields, such as machine learning, credit scoring, medical research, and polygenic risk score prediction, where binary classification is important. It provides a measure of a predictive model’s ability to distinguish between positive and negative instances. The calculation of AUC is based on the receiver operating characteristic (ROC) curve, which plots the true positive rate (sensitivity) against the false positive rate (1 - specificity) at different classification thresholds. The AUC represents the area under the ROC curve and ranges from 0 to 1. A perfect classifier has an AUC of 1, indicating flawless discrimination, while an AUC of 0.50 suggests a classifier that performs no better than random chance. Generally, a higher AUC value indicates better discrimination power and a more accurate model.

To construct the empirical ROC curve, consider a sample of individuals consisting of *m* cases and *n* controls. The predictor variable is continuous, and higher values are assumed to be associated with the cases. The prediction performance can be assessed by calculating the sensitivity and specificity at various threshold values (*v*). Sensitivity (*v*) is the proportion of cases with values above the threshold, and specificity (*v*) is the proportion of controls with values below the threshold. Mathematically, they can be expressed as:

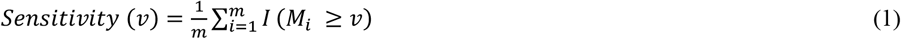

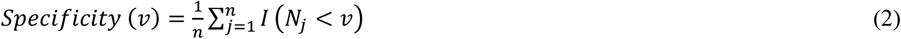

where *M*_*i*_ and *N*_*j*_ represent the values of the predictor variable for the *i*^*th*^ individual in the case group and the *j*^*th*^ individual in the control group, respectively. The indicator function *I*(A) equals 1 if the condition A is true and 0 otherwise.

By varying the threshold value (*v*) over the range of possible values, we can plot the empirical ROC curve. The curve starts at the point (0, 0) when the threshold is higher than the largest possible value and gradually increases towards the point (1, 1) as the threshold decreases to the smallest possible value. Ideally, the entire curve should lie above the diagonal line where sensitivity equals 1 – specificity, indicating better-than-chance classification performance.

Furthermore, according to DeLong et al.^16^, this AUC is equivalent to the Mann-Whitney two sample statistic applied to the two samples *M*_*i*_ and *N*_*j*_. The Mann-Whitney statistic, denoted as ***ω***, can be expressed as:

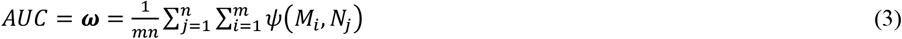

In this equation, *ψ*(*M*_*i*_, *N*_*j*_) equals 1 if *N* < *M*, ½ if *N* = *M*, and 0 if *N* > *M*.

Another equivalent definition of AUC is the probability that a randomly selected pair of case and control are accurately classified. Wray et. al.^11^ suggested that this probability can be approximated as the probability that difference between the predicted values of the cases and controls is greater than zero. The predicted values refer to a single set of PRS or a composite vector derived from multiple sets of PRSs in this context. The difference in predicted values follows an approximate normal distribution with a mean of 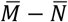 and a variance of 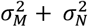, as proposed by Wray et al.^11^. Following Wray et al.^11^ and the truncated normal distribution theory, the mean and variance of the difference can be calculated as

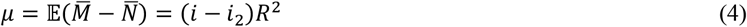

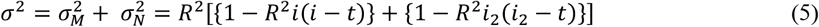

with 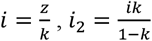,
where *R*^2^ represents the coefficient of determination in a linear model with the liability, which denotes the proportion of variance explained by the PRS on the liability scale. Additionally, *k* represents the population lifetime prevalence, *z* is the height of a normal density curve at a specific point according to *k*, and t is the threshold that truncates the normal distribution according to *k*. The mean liability for cases is denoted as *i*, and the mean liability for controls is represented as *i*_2_.

The predicted values of the cases and controls are typically assumed to follow a normal distribution, unless the *R*^2^ value is exceptionally high^11^ (**Figure S1**).

Therefore, the expected AUC can be calculated using the cumulative density function (Φ) with the formula:

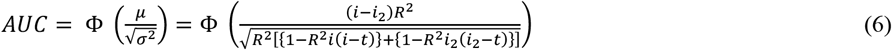

Conversely, given an AUC value, *R*^2^ can be derived using the equation

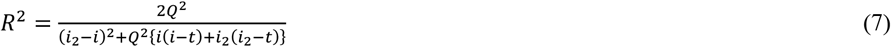

where Q represents the Φ^−1^(*AUC*) (Wray et al.^11^)

Additionally, it is important to note that *R*^2^ values on the liability scale and the observed scale can be transformed to each other^19^. For example, the transformation equation for a population cohort is as follows:

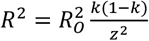

where *R*^2^ is the coefficient of determination on the liability scale, 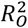 denotes the coefficient of determination on the observed scale, which is applicable for binary disease outcomes, and *k* and *z* were defined above. This transformation is particularly useful when working with binary responses and allows for the conversion between functions of 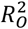 and AUC.

### The DeLong test to compare AUCs of two Models

The DeLong test^16^ is a statistical method commonly used to compare the performance of two models based on their AUC values. It is employed to assess whether there is a statistically significant difference in the predictive ability of the models. To conduct the DeLong test, AUC estimates and their corresponding variances are required for both models. These estimates can be obtained through various techniques, such as using receiver ROC analysis or fitting predictive models. The DeLong test calculates the covariance matrix between the AUC estimates of the two models. It estimates the covariance, taking into account the correlation between the AUC estimates, and constructs a test statistic. This nonparametric approach, known as DeLong’s method, is widely used in statistical analysis for comparing AUCs without making assumptions about the underlying data distribution. It provides a robust estimate of the covariance term, accounting for the variability in the false positive rate (FPR) between the models. By considering the covariance term, DeLong’s method enables an accurate assessment of the statistical significance of the difference in AUCs.

DeLong’s method, using the Heaviside function, can be computationally demanding. However, Sun et al.^17^ proposed an efficient approach that estimates the variance and covariance matrix between two AUC models by leveraging the relationship between the Heaviside function and the mid-ranks of the samples. This improvement reduces the computational burden from quadratic to linearithmic. The DeLong test statistic follows a normal distribution, allowing for the calculation of a p-value. The p-value indicates the significance of the observed difference in AUCs between the models, providing valuable insights for model comparison. For more detailed information on the variance and covariance matrix estimation, please refer to DeLong et al.^16^ and Sun et al.^17^.

### Proposed method to compare AUCs of two models

To compare the AUC values of two models, we employ the delta method to estimate the variance of the difference between the AUC values based on two sets of PRS denoted as x_1_ and x_2_, with respect to the underlying phenotypes y. The difference in AUC values can be formulated as a function of sets of correlations, denoted as 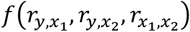. It is noted that one AUC is a function of the correlation 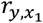, while the other AUC is a function of the correlation 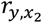, as shown in equation (6). The delta method approximates the variance of the difference between the two AUC values as

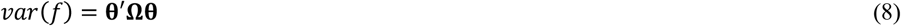

where 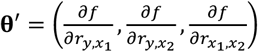 is the derivatives of *f* with respect to the correlations
and

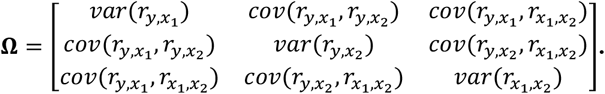

Each element of **Ω** represents the sampling variance of the correlation, as shown in Olkin and Finn^20^ and Momin et al.^21^. From eq. (8), the variances of differences can be estimated and used in our analyses of AUC based on PRS.

### Assessing AUC difference in PRS models generated from different GWAS discovery datasets (i.e. non-nested models)

Using eq. (6), the variance of AUC difference can be written as

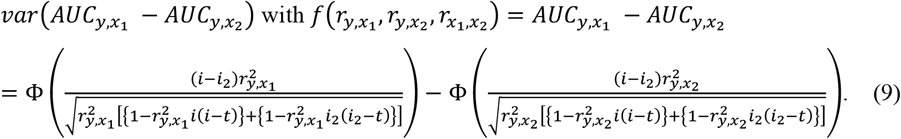

The variance of the AUC difference can be obtained using eq. (8) with the derivatives with respect to each of the correlations and the variance and covariance matrix of the correlations. In eq. (9), the values of 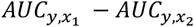 from random samples in the population are assumed to follow a normal distribution^16, 17^. Therefore, the p value for the significance test of the difference can be derived from 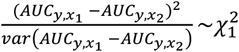.

Additionally, the 95% confidence interval for the difference in AUC values can be calculated as

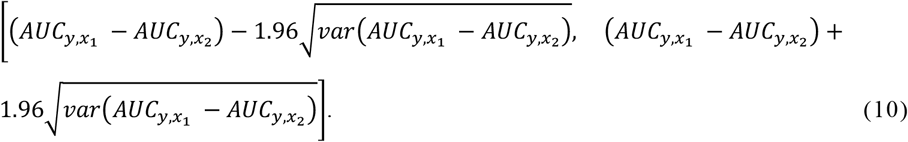

By applying this proposed method, we can efficiently compare the AUC models with different PRS predictors (*x*_1_ and *x*_2_) that may or may not be correlated with each other. It provides a statistical framework to assess the significance of the difference between the AUC values and estimate confidence intervals for the comparison.

### Assessing AUC difference in nested PRS models

When comparing nested models, the variance of the AUC difference can be expressed as follows:

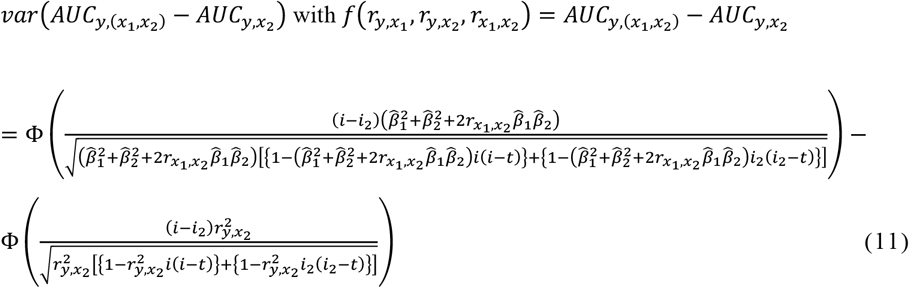

where 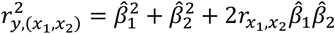 and 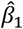 and 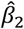 are the estimated regression coefficients from a multiple regression^21^ as

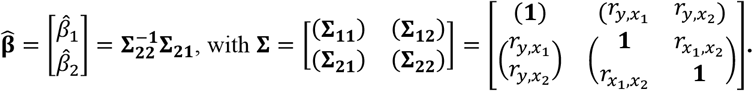

Therefore, eq. (11) can be expressed as a function of the correlations, 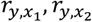 and 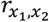, and the derivative with respect to each of the correlations can be obtained for this function.

Following DeLong et al.^16^, it is assumed that the values of 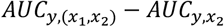 in eq. (11) follow a normal distribution when *x*_1_ is associated with the outcome^22^. Consequently, the p value for the significance test of the difference can be derived from 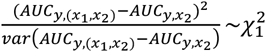.

Furthermore, the 95% confidence interval can be calculated as follows:

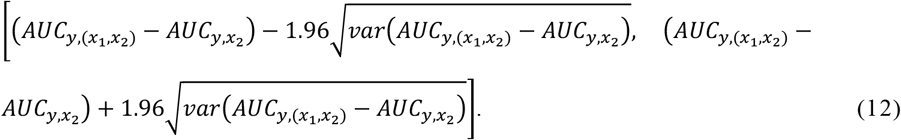

However, it is crucial to acknowledge that if *x*_1_ is not significantly associated with the outcome, the DeLong’s test may result in a conservative test because AUC difference between the nested models does not follow a normal distribution, violating the assumption made in the DeLong’s test^22^. In such cases, alternative approaches based on likelihood ratio test^23^ or *R*^2^-based test^21^ can be employed for comparing nested models. For further information on the DeLong’s test and its application in nested model comparison, please refer to additional references^22, 24^.

### Assessing AUC difference when using two independent sets of PRS

It is of interest to use two independent sets of PRS for two distinct sets of target samples, such as PRS calculated separately for male and female individuals or in non-overlapping cohorts. In such a comparison, it is worth noting that one can compare their estimated AUC to a published AUC, provided that the variance (or 95% confidence interval) of AUC is known. In these scenarios, there is no correlation structure between two independent sets of PRS 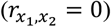. Hence, the variance of AUC difference is simply the sum of the variances of each AUC value, which can be obtained^16, 25^. For example, assuming 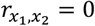, the variance of *R*^2^ difference can be written as

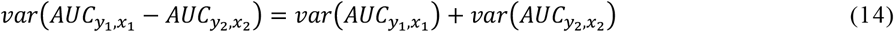

where,

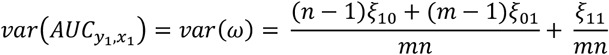

with

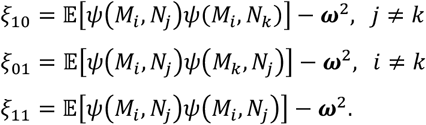

Alternatively, using eq. (8), this variance can also be efficiently derived as

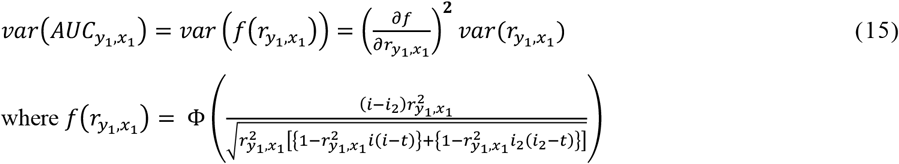

In a similar manner, 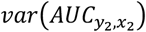 can be obtained and eq. (14) can be solved.

The p-value for the significance test of the difference can be derived from 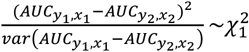 and the 95% confidence interval^20^ is calculated as

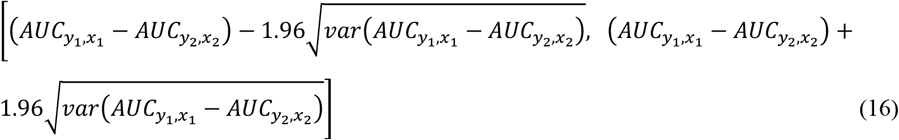

### Simulation of binary outcomes using a liability threshold model

Simulation of binary outcomes using a liability threshold model is a common and useful approach in statistical genetics and epidemiology, particularly for method verification purposes. The liability threshold model assumes that an individual’s liability to a particular binary trait is normally distributed in the population. If the liability surpasses a certain threshold, the individual is considered affected; otherwise, they are unaffected. To simulate binary outcomes using a liability threshold model, we used the following steps. Firstly, we set the parameters for the simulation, including the mean and standard deviation of liability distribution in the population (0 and 1, respectively). We also set the threshold value that determines the affected/unaffected status. We explored various simulation scenarios by using multiple threshold values (e.g., *t* = 1.64, 1.28, 1.04, 0.84, and 0), resulting in various disease prevalence (e.g., *k* = 0.05, 0.10, 0.15, 0.2, and 0.5). Next, we generated the dependent variable, y, representing the liability to the trait for each individual, as well as PRSs (*x*_1_ and *x*_2_), using a multivariate normal distribution. The correlation structure was represented by the matrix 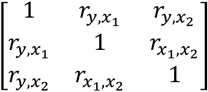, which varied according to simulation scenarios. Then, we assigned affected or unaffected status according to the liability values, compared to the threshold. The transformed affected/unaffected status were used in the analyses while continuous *x*_1_ and *x*_2_ were used as they were. This simulation approach allows us to generate binary outcomes based on a specified liability threshold, reflecting the underlying distribution of liability in the population. It enables us to explore various scenarios and comprehensively evaluate R2ROC and DeLongs’ approaches.

### Real data

To apply the proposed method, R2ROC, to a real dataset, we used the UKBB. Detailed information about the dataset can be found in elsewhere^26, 27^. Briefly, the UKBB is a comprehensive biomedical database comprising 500,000 volunteers recruited between 2006 and 2010, aged between 40 and 69 years^26, 28^. It contains extensive health-related information and genome-wide SNP genotyping data. For our analysis, we utilized the genotype data that passed a rigorous quality control (QC) process as described in Momin et al.^21^. Following QC, the dataset consisted of 288,792 unrelated white British individuals (genomic relationship matrix coefficient <0.05), based on their principal component (PC) and 7,701,772 SNPs. In this analysis, we included individuals with type 2 diabetes (T2D) diagnosed by a doctor and coronary artery disease (CAD) based on ICD-10 information. Among the total of 288,792 individuals, there were 13,772 cases of T2D and 27,898 cases of CAD, while the remaining individuals served as controls.

To evaluate AUC values, we divided the 288,792 individuals into discovery and target datasets by randomly selecting 90% and 10% of the individuals, respectively. This resulted in a discovery dataset of 259,912 individuals and a target dataset of 28,880 individuals. In the discovery dataset, we conducted a genome-wide association study (GWAS) analysis on diabetes and CAD, controlling potential confounding factors such as age, sex, birth year, Townsend deprivation Index (TDI), education, genotype measurement batch, assessment centre, and the first 10 ancestry principal components.

In addition, we have access to GWAS summary statistics from the Japanese Biobank (BBJ) for diabetes and CAD. The diabetes dataset comprises 36,614 cases and 155,150 controls, encompassing a total of 12,557,762 single nucleotide polymorphisms (SNPs) for analysis^29^. On the other hand, the CAD dataset includes 29,319 cases and 183,134 controls, with 6,046,444 SNPs available for investigation^30^.

Based on these two sets of UKBB and BBJ GWAS summary statistics, we constructed two sets of PRS for the 28,880 individuals in the target dataset for each disease. This estimation was performed using the “*--score*” function in PLINK2 software^31^. Ambiguous SNPs and SNPs with any strand issues were excluded from the analysis, resulting in the use of 4,067,234 SNPs for diabetes and 3,644,268 SNPs for CAD that were common between the UKBB and BBJ GWAS datasets.

We applied the R2ROC method to the target data, fitting the constructed PRS and binary outcomes (diabetes and CAD), to estimate the AUC and evaluate the AUC difference between the PRS derived from UKBB and BBJ. Within the target dataset consisting of 28,880 individuals, we simultaneously modelled the phenotypes for each trait alongside covariates such as age, sex, birth year, Townsend deprivation Index (TDI), education, genotype batch, assessment centre, and the first 10 principal components. These covariates were incorporated to account for potential confounding factors and ensure accurate modelling of the PRS.

## Results

### Simulation verification

The performance of the proposed method has been examined through comprehensive simulations. The simulations involved varying the correlation structure of 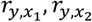 and 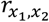, as well as the disease prevalence rate (*k*) (see simulation section in the Methods). These variations allowed a thorough comparison between non-nested PRS models (i.e., 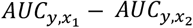) and nested PRS models *(i*.*e*., 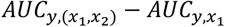and 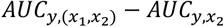). The performance evaluations were conducted based on 10,000 simulated replicates, from which the AUC differences and variances of the differences were derived for both DeLong’s method and the proposed R2ROC method. Notably, an almost perfect correlation was observed between DeLong’s and R2ROC methods for both the AUC differences and variances of the differences whether using non-nested or nested PRS models when considering a realistic correlation structure between 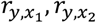 and 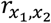 (**Figure 1**). When varying the correlation structure, e.g. 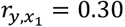, 0.45, and 0.60, we consistently observed that DeLong’s and R2ROC methods generated very similar results (correlation > 0.999) (**Figures S2-S4**). **Table S1-S3** demonstrate that the mean differences of AUC and their variances, obtained from 10,000 simulated replicates, are similar between the DeLong’s method and R2ROC method. This result demonstrates the robustness and reliability of the proposed method, affirming its performance in accurately assessing AUC differences across different correlation structures and disease prevalence levels.

**Figure 1:**
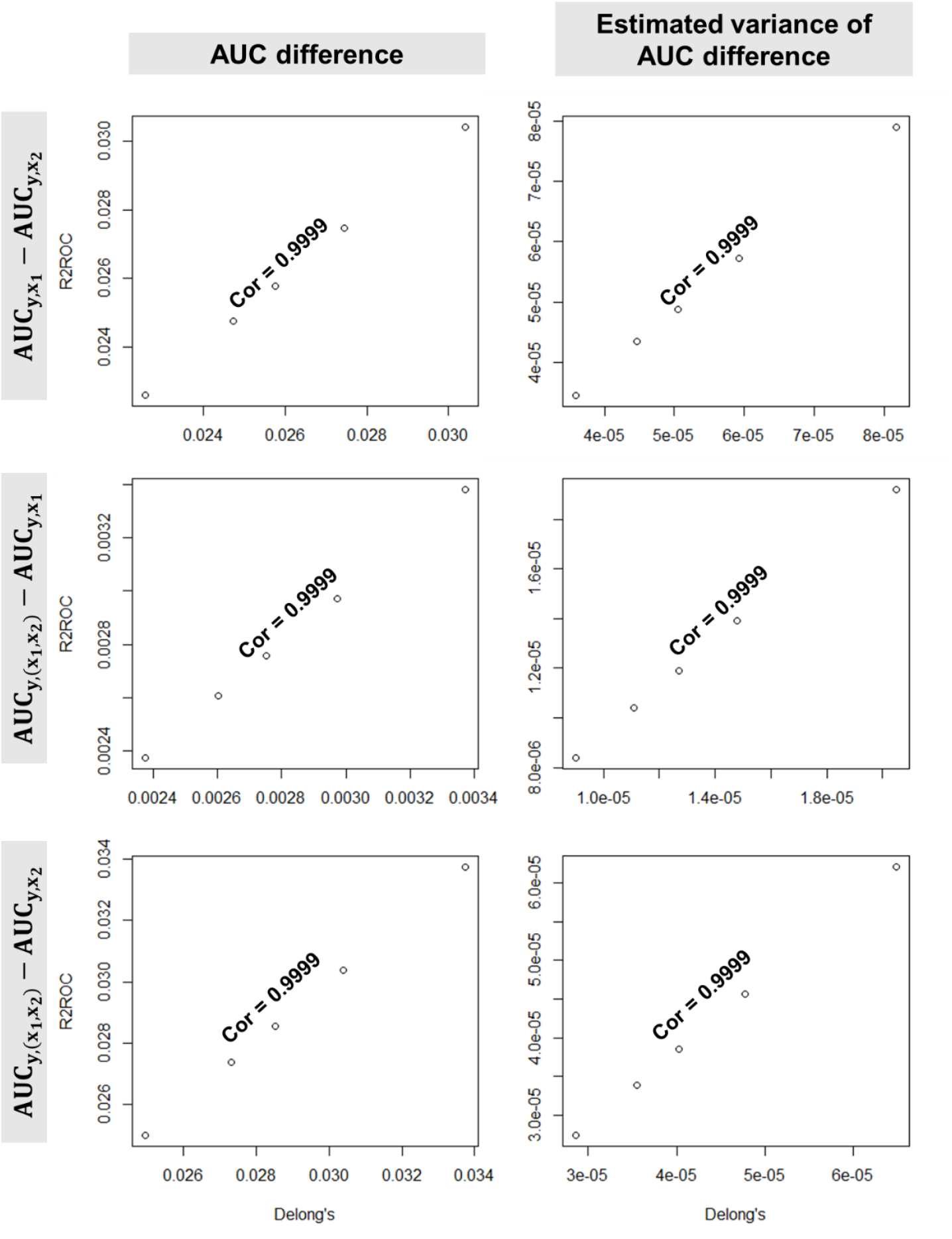
Agreement between DeLong’s and R2ROC methods for AUC difference and estimated variance of AUC difference when varying population prevalence levels. The dots represent the averages of AUC differences and estimated variances of AUC difference calculated from 10,000 simulated replicates, revealing the strong agreement between DeLong’s and R2ROC methods across various population prevalence levels (*k*=0.05, 0.10, 0.15, 0.20, and 0.50). The comparison involves both non-nested 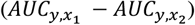 and nested PRS models 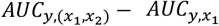 and 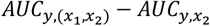. To generate the dependent variable (y) representing liability values, as well as the PRSs (*x*_1_ and *x*_2_), a multivariate normal distribution was employed. The correlation structure was determined by the matrix 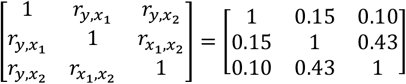, mimicking the real correlation structure between UKBB and BBJ PRSs for diabetes. Each simulation involved 30,000 individuals to ensure robustness and statistical power.

### Computational efficiency of R2ROC

The accuracy and reliability of the R2ROC method are evident in its performance, i.e. the R2ROC method exhibits comparable precision to DeLong’s test in estimating differences between two AUC values and their variances, as demonstrated in **Figure 1** and **Figures S2-S4**. However, the R2ROC method surpasses DeLong’s method in terms of computational efficiency, as shown in **Figure 2**. The computational speed of the R2ROC method is up to 150 times faster than that of DeLong’s method. For example, when analysing data from 5,000 individuals, DeLong’s method took over 70 times longer to compute compared to R2ROC in estimating the AUC difference and the variance of AUC difference. When analyzing a larger sample size of 400,000, the computational efficiency of R2ROC was substantially higher than DeLong’s method, with around 150-fold difference in speed. It is worth noting that the most computationally optimized version of DeLong’s method, implemented using the pROC package^18^ with the latest version 1.18.4, was used in the analysis to ensure a fair comparison^18^.

**Figure 2:**
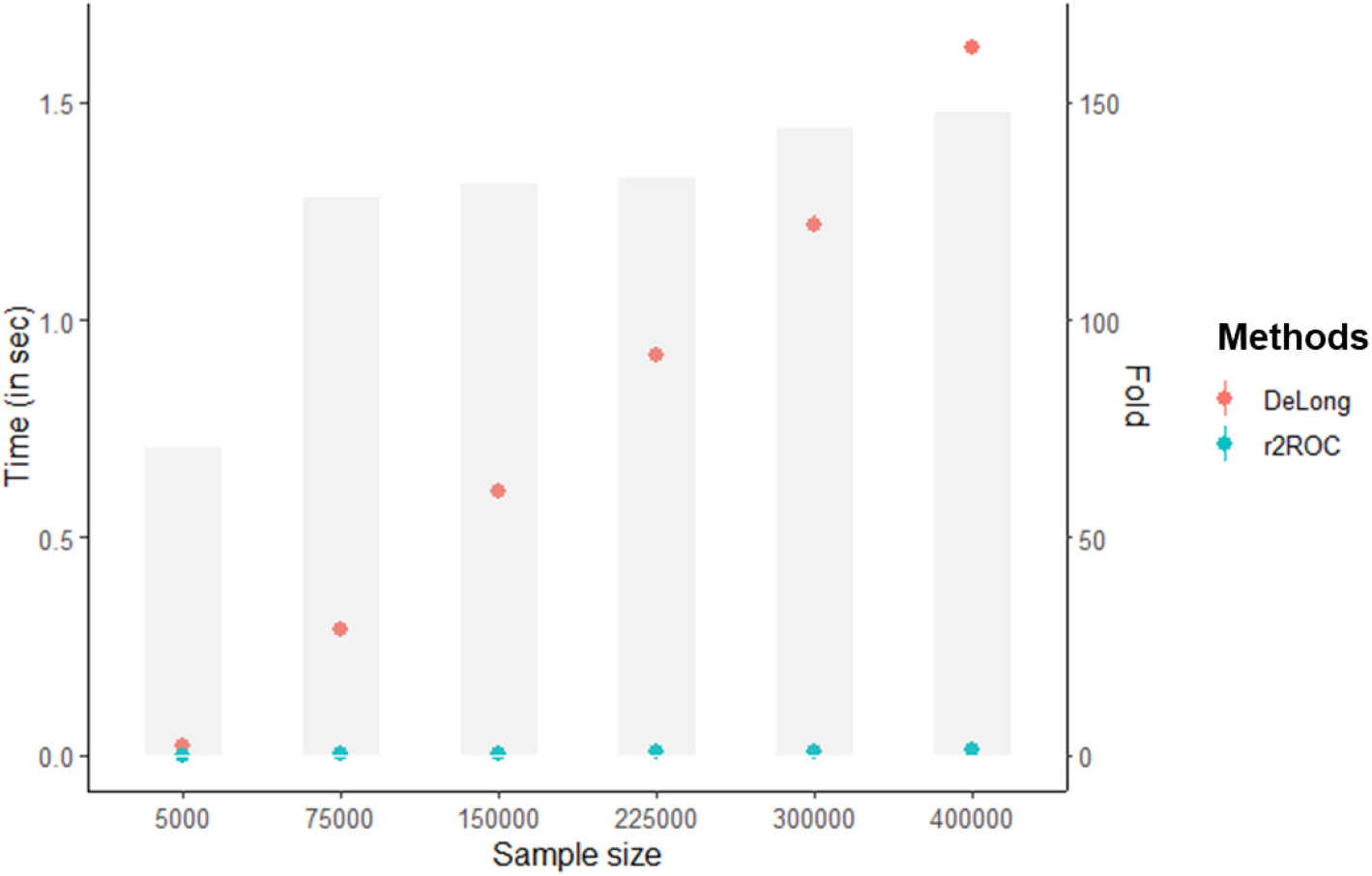
Comparison of computing time between DeLong’s and R2ROC methods with varying sample sizes. The dot points represent the time taken by each method to estimate the AUC difference and the variance of AUC difference, while the bar indicates the fold-difference in speed, illustrating the relative efficiency of the R2ROC method compared to DeLong’s method. The measured CPU times exhibited little variation across replicates, allowing us to conduct a small number of replicates (n=10) to obtain the average with negligible sampling variance.

### Comparative performance evaluation of PRS Models for diabetes and CAD using R2ROC Method

In this section, we performed a comparative performance evaluation of two sets of PRS models for diabetes and CAD using the R2ROC method. The first set of PRS models was constructed using the UKBB discovery dataset (referred to as UKBB PRS), while the second set utilized the BBJ discovery dataset (referred to as BBJ PRS). The AUC analysis was conducted on the independent target cohort comprising 28,880 individuals with white British ancestry.

**Figure 3** presents the AUC values for diabetes, which were estimated to be 0.593 (95% CI = 0.573 - 0.608) for the UKBB PRS and 0.555 (95% CI = 0.570 - 0.539) for the BBJ PRS. Likewise, the AUC values for CAD were 0.595 (0.584 - 0.606) and 0.575 (0.564 - 0.586) for UKBB and BBJ PRS, respectively. However, these AUC values and confidence intervals cannot be used to assess the difference between the two sets of PRS due to their lack of independence. Additionally, the UKBB and BBJ PRS models are not nested, making the likelihood ratio test (LRT) unsuitable for comparison. The DeLong method, although applicable, is known to be computationally demanding^16, 17^.

**Figure 3:**
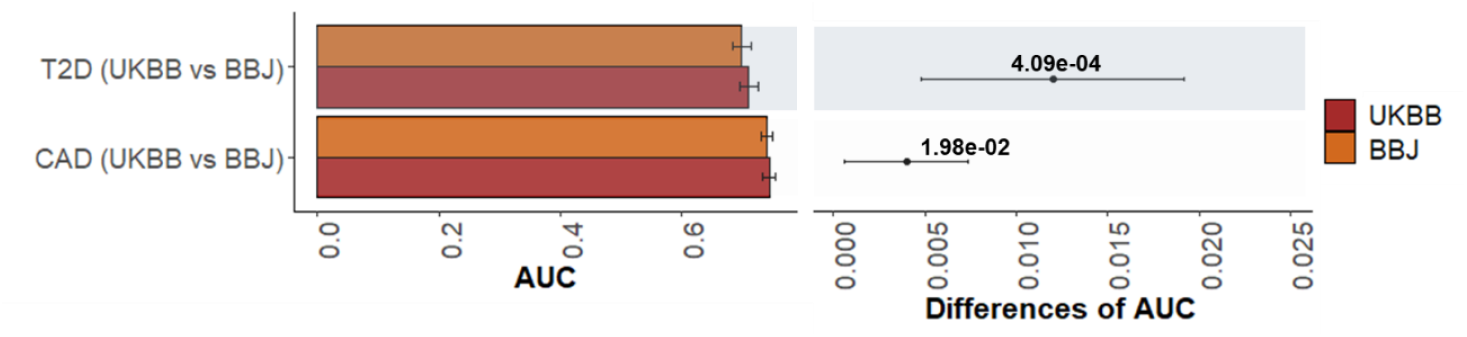
Comparative performance evaluation of PRS models for diabetes and CAD (N=28,880) using PRS derived from UKBB and BBJ. **Left panel:** The main bars represent AUC values and error bars correspond 95% confidence intervals. Two sets of GWAS summary statistics were obtained from UKBB and BBJ discovery GWAS datasets to estimate two sets of PRS in the target dataset of 28,880 European individuals (UKBB PRS vs. BBJ PRS). **Right panel:** Dot points represent the differences of AUC values between UKBB and BBJ PRS models, and error bars indicate 95% confidence intervals of the difference. P-values are provided to indicate the significance of the difference between the compared pairs of PRS models (UKBB PRS vs. BBJ PRS). In the PRS models, we simultaneously modelled the phenotypes for each trait alongside covariates such as age, sex, birth year, Townsend deprivation Index (TDI), education, genotype batch, assessment centre, and the first 10 principal components.

To overcome these challenges, we employed the R2ROC method (implemented in R2ROC package) for a comparative significance test between two sets of AUC values (from UKBB PRS vs. BBJ PRS models) for diabetes and CAD. The calculated difference in AUCs for diabetes was 0.012 (95% CI = 0.0196 - 0.0056) with a p-value of 4.09e-04 **(Figure 3)**. Similarly, for CAD, the AUC difference had a 95% confidence interval of 0.0006 to 0.007 and a p-value of 1.98e-02 (**Figure 3**). This comparative performance evaluation demonstrates the effectiveness of the R2ROC method in accurately assessing the differences between non-nested PRS models for diabetes and CAD. By using R2ROC method, we overcame the challenges associated with non-nested models and obtained reliable estimates of the AUC differences specific to the PRS of interest, along with their statistical significance.

While non-nested model comparison provides a valuable approach to evaluating predictors (as shown in **Figure 3**), nested model comparison offers an additional perspective by assessing whether the inclusion of additional predictors or parameters significantly improves the fit of a more complex model compared to a simpler model. In the context of diabetes and CAD data, it is important to determine if the BBJ PRS, as an additional predictor, enhances the AUC significantly when added to the UKBB PRS, or vice versa.

**Figure 4** presents a comparison of the AUC values obtained from each UKBB or BBJ PRS individually, as well as from a joint model that incorporates both UKBB and BBJ PRS simultaneously. The AUC difference observed between the joint model and the individual model using only the UKBB PRS demonstrates a statistically significant improvement (p-value 6.64e-03) in diabetes prediction with the inclusion of the BBJ PRS (see **Figure 4**). Similarly, the test statistics for the AUC difference between the joint model and the model using only the UKBB PRS indicate significance for CAD, with a 95% confidence interval of 0.0016 to 0.0049 and a p-value of 1.53e-04 (see **Figure 4**). This analysis provides evidence for the added value of incorporating both PRS datasets in predicting diabetes and CAD, beyond the use of either PRS individually.

**Figure 4:**
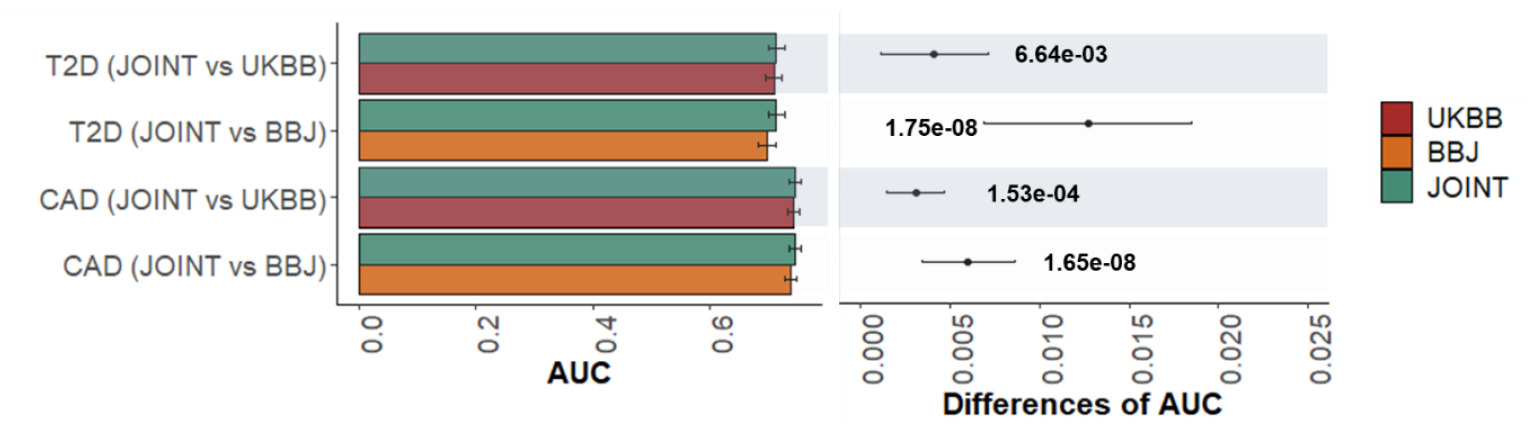
Comparative performance evaluation of nested models for diabetes and CAD. **Left panel:** The main bars represent AUC values and error bars correspond 95% confidence intervals. Two sets of GWAS summary statistics were obtained from UKBB and BBJ discovery GWAS datasets to estimate two sets of PRS, i.e., UKBB and BBJ PRS. In addition, a joint model fitting both UKBB and BBJ PRS was compared. **Right panel**: Dot points represent the differences of AUC values between the joint model and UKBB or BBJ PRS model, and error bars indicate 95% confidence intervals of the difference. P-values are provided to indicate the significance of the difference between the compared pairs. In the PRS models, we simultaneously modelled the phenotypes for each trait alongside covariates such as age, sex, birth year, Townsend deprivation Index (TDI), education, genotype batch, assessment centre, and the first 10 principal components.

## Discussion

Genetic analysis using polygenic risk scores (PRSs) has emerged as a promising approach for early prediction of the genetic contribution to disease risk^9^. In this study, we focused on evaluating the performance of PRSs for binary disease traits using the AUC. While AUC is widely used for assessing PRS performance, the commonly employed DeLong’s test for comparing PRS by comparing AUC has certain limitations^22, 32^ that prompted the development of a novel approach using the Delta method. Our proposed approach addresses several limitations of DeLong’s test and offers significant advantages.

Firstly, it reduces computation time by up to 150-fold, enabling efficient large-scale analyses while generating identical results. This improvement is crucial as genetic studies often involve extensive datasets and require computationally efficient methods for practical implementation. The reduced computational burden facilitates the application of our approach to real-world datasets, enhancing its practicality and utility. This can be also useful in a machine learning where repeated comparative significance tests are required^33^. Importantly, considering the rising concern to minimize greenhouse gas emissions^34^, R2ROC emerges as the more environmentally friendly option, emitting significantly lower carbon emissions during computation. For instance, in iterative analyses of 100,000 comparative significance tests using data from 400,000 participants, DeLong’s test demands an average of approximately 45 hours, leading to a substantial carbon footprint of about 300 kg CO_2_ according to the carbon footprint calculator (http://calculator.green-algorithms.org/)^34^. In contrast, R2ROC completes the same analyses in a mere 0.5 hours, resulting in a significantly lower carbon footprint of approximately 3 kg CO_2_.

Another significant aspect of our study is the exploration of the transferability between R^2^ and AUC. We established a relationship and one-to-one mapping between R^2^ and AUC in terms of comparative significance tests. This equivalency provides valuable insights into how changes in model performance, as measured by R^2^, correspond to changes in discriminative ability, as measured by AUC. This understanding aids in interpretation, assists in model selection or comparison, and sheds light on the generalizability of model performance across these two metrics.

Additionally, the established relationship between the test statistics of AUC and R^2^ allows for easy interpretation of individual variable effects, providing valuable insights into the specific contributions of variables to overall AUC values. This interpretability is particularly important when multiple variables are involved in AUC computation, as it enhances our understanding of the genetic factors influencing disease risk. By gaining insights into the relative importance of these variables, researchers and clinicians can prioritize interventions more effectively, leading to improved personalized disease prediction and informed healthcare decision-making.

To validate the effectiveness of our proposed approach, comprehensive simulations were conducted, demonstrating its robustness and reliability. Furthermore, we applied our approach to real data from a cohort of 28,880 European individuals to compare PRSs for diabetes and coronary artery disease (CAD) prediction. The use of genome-wide association study summary statistics from two distinct sources allowed us to comprehensively assess the predictive abilities of the PRSs. Through this analysis, we shed light on the respective strengths and weaknesses of the PRSs in predicting diabetes and CAD, contributing to the assessment of genetic risk factors associated with these diseases.

There are several limitations in this study. Firstly, when the R^2^ value from a prediction model is very high (>0.5), the approximation becomes poor. However, it is important to note that such high R^2^ values are exceptional in the PRS context. Secondly, we did not investigate the relationship between the likelihood ratio test and R2ROC test for nested model comparisons, as well as the relationship between Vuong’s test and R2ROC test for non-nested model comparisons. It is worth mentioning that Vuong’s test is a well-known likelihood ratio test specialized for non-nested model comparisons. Investigating these relationships could provide valuable insights into their comparative performance for model evaluations, which is another avenue to be explored in future research. Moreover, we did not explicitly test a model using pre-adjusted outcomes, which is commonly used in the genetic field. This is an important consideration that warrants investigation in future studies to understand how pre-adjusted outcomes impact PRS performance and model comparisons in such comparative significance tests of AUC.

In summary, our novel approach utilizing the Delta method overcomes limitations of the widely used DeLong’s test and offers significant advantages, including reduced computation time and enhanced interpretability of individual variable effects. By applying our approach to real data, we demonstrate its effectiveness in assessing the predictive abilities of PRSs for diabetes and CAD. This advancement in genetic risk assessment and personalized disease prediction has the potential to greatly impact healthcare decision-making, ultimately leading to improved patient outcomes.

## Data and Code availability

The genotype and phenotype data of the UK Biobank can be accessed by following the procedures described on its webpage (https://www.ukbiobank.ac.uk/). On this website, you will find information on how to access and utilize the data for research purposes. For summary statistics of diabetes and CAD from the Japanese Biobank (BBJ) and could have obtained them from the BBJ website (http://jenger.riken.jp/en/).

pROC can be downloaded from (https://github.com/xrobin/pROC) or from CRAN.

R2ROC can be downloaded from (https://github.com/mommy003/R2ROC) or from CRAN.

## Acknowledgements

This research has received support from the Australian Research Council (DP190100766). We would like to express our gratitude to the staff and participants of the UK Biobank and Biobank Japan for their valuable contributions to this study. Our research project has been assigned reference number 14575 by the UK Biobank. We would also like to acknowledge the computational resources provided by the Australian Government through Gadi under the National Computational Merit Allocation Scheme (NCMAS). Additionally, we would like to thank UniSA IT for managing the HPCs (StatGen and StatGen2 server) that were utilized for the data analyses in this research. NRW acknowledges funding from the National Health and Medical Research Council (1173790).

## Author contribution

S.H.L. and N.R.W. conceived the idea. S.H.L. derived theory and supervised the study. M.M.M performed the analysis. M.M.M and S.H.L made the R package. S.H.L and M.M.M wrote the first draft of the manuscript. N.R.W. provided critical feedback and suggestions. All the authors discussed the results and contributed to the final manuscript.

