## Supplementary material for "R2ROC: An efficient method of comparing two or more correlated AUC from out-of-sample prediction using polygenic scores": AUC_manuscript_Supplementary_02.08.2023_final_submission.pdf

### Supplemental figures

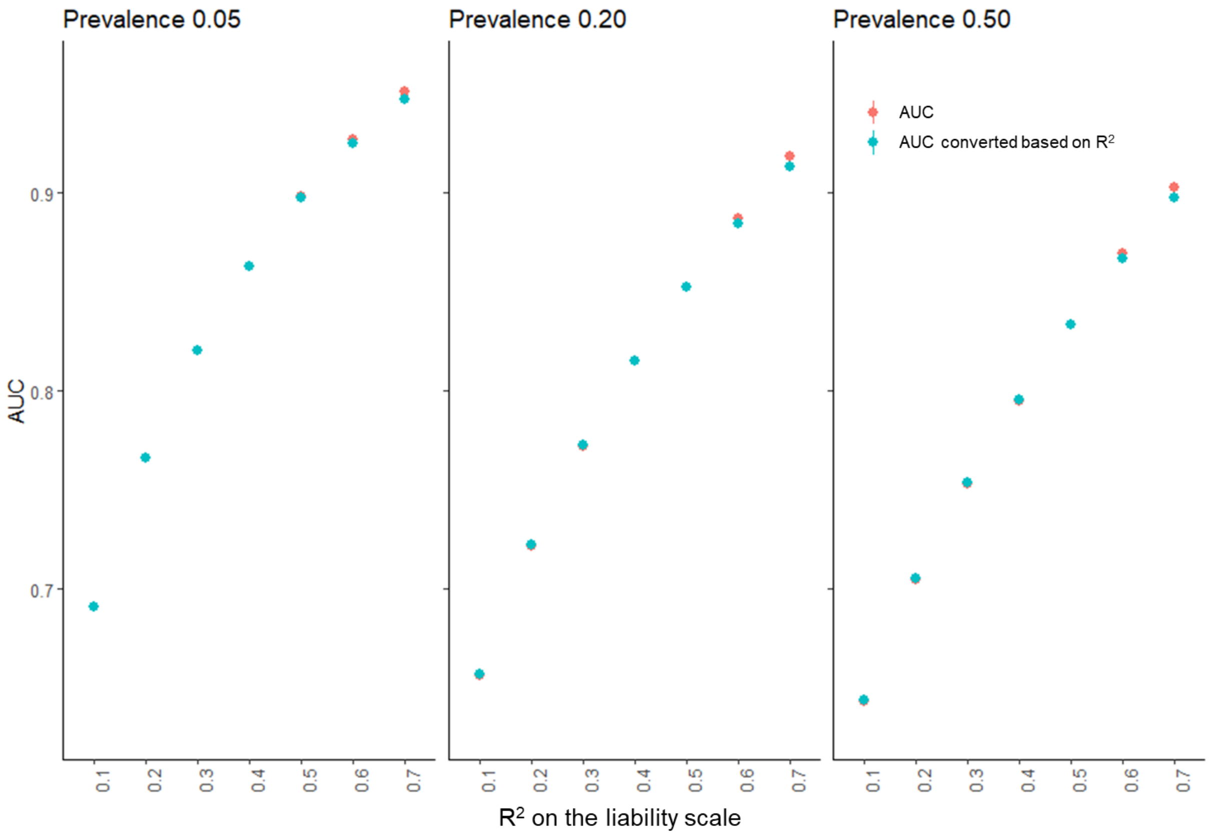


**Figure S1: AUC can be accurately estimated based on $R^2$ unless $R^2$ value is exceptionally high.** $R^2$ value on the liability scale resulting from the association between disease outcome and PRS predictor typically ranges from 0.02 to 0.25[1-5], indicating that AUC can be accurately estimated based on $R^2$ in general. The dots represent the averages of AUC estimates from 1,000 simulated replicates using various population prevalence levels (*k*=0.05, 0.20, and 0.50). To generate the dependent variable (y) representing liability values, as well as the PRS ($x_1$), a multivariate normal distribution was employed. The correlation structure was determined by the matrix $\begin{bmatrix} 1 & r_{y,x_1} \\ r_{y,x_1} & 1 \end{bmatrix} = \begin{bmatrix} 1 & various \\ various & 1 \end{bmatrix}$. The correlation structure was changed based on $R^2$ values on the liability scale. Each simulation involved 30,000 individuals.

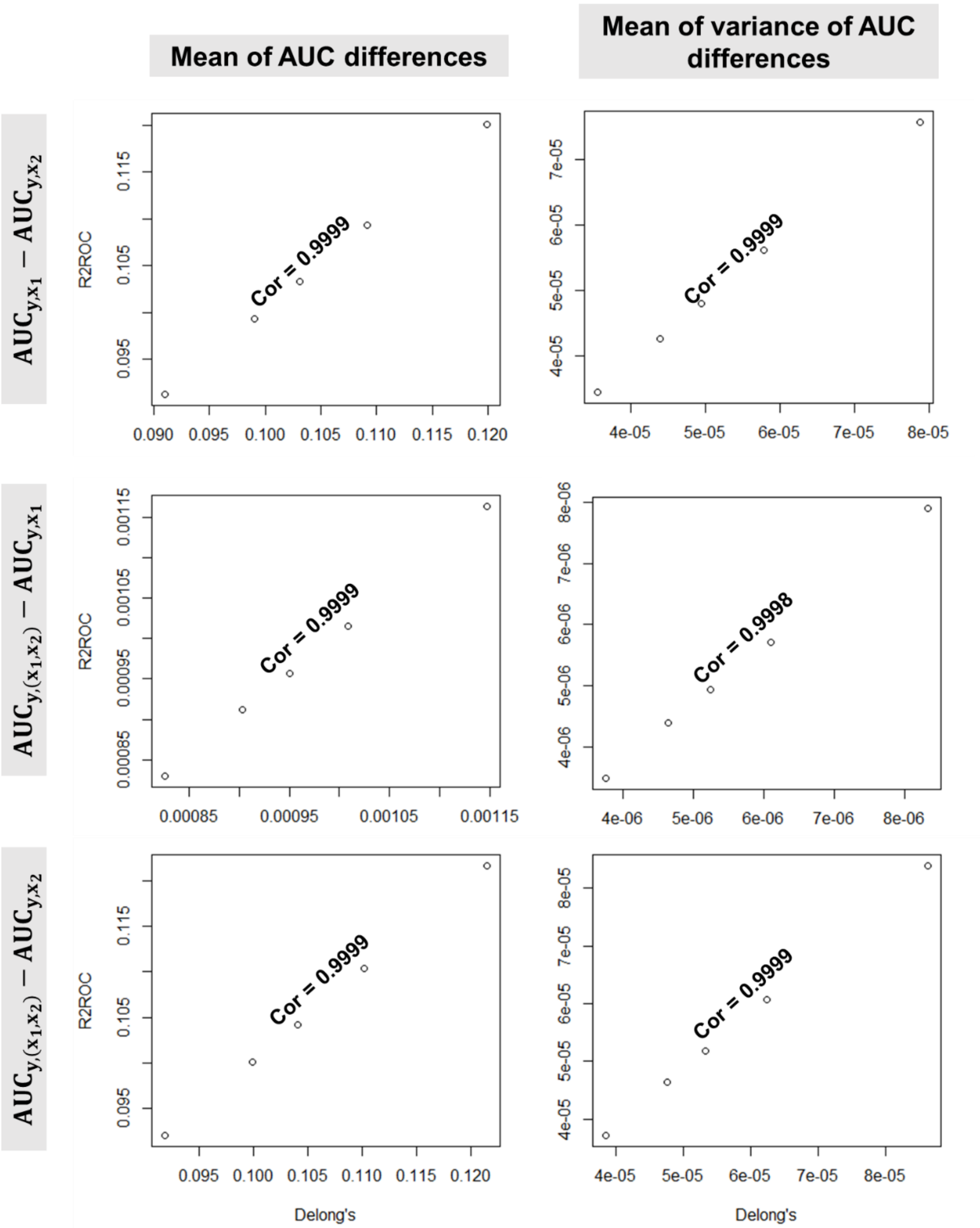


**Figure S2: Agreement between DeLong's and R2ROC methods for AUC difference and estimated variance of AUC difference when varying population prevalence levels.**

The dots represent the averages of AUC differences and estimated variances of AUC difference calculated from 10,000 simulated replicates, revealing the strong agreement between DeLong's and R2ROC methods across various population prevalence levels (*k*=0.05, 0.10, 0.15, 0.20, and 0.50). The comparison involves both non-nested ($AUC_{y,x_1} - AUC_{y,x_2}$) and nested PRS models ($AUC_{y,(x_1,x_2)} - AUC_{y,x_1}$ and $AUC_{y,(x_1,x_2)} - AUC_{y,x_2}$). To generate the dependent variable (y) representing liability values, as well as the PRSs ($x_1$ and $x_2$), a multivariate normal distribution was employed. The correlation structure was determined by the matrix $\begin{bmatrix} 1 & r_{y,x_1} & r_{y,x_2} \\ r_{y,x_1} & 1 & r_{x_1,x_2} \\ r_{y,x_2} & r_{x_1,x_2} & 1 \end{bmatrix} = \begin{bmatrix} 1 & 0.30 & 0.10 \\ 0.30 & 1 & 0.43 \\ 0.10 & 0.43 & 1 \end{bmatrix}$, mimicking the real correlation structure between UKBB and BBJ PRSs for diabetes. Each simulation involved 30,000 individuals to ensure robustness and statistical power.

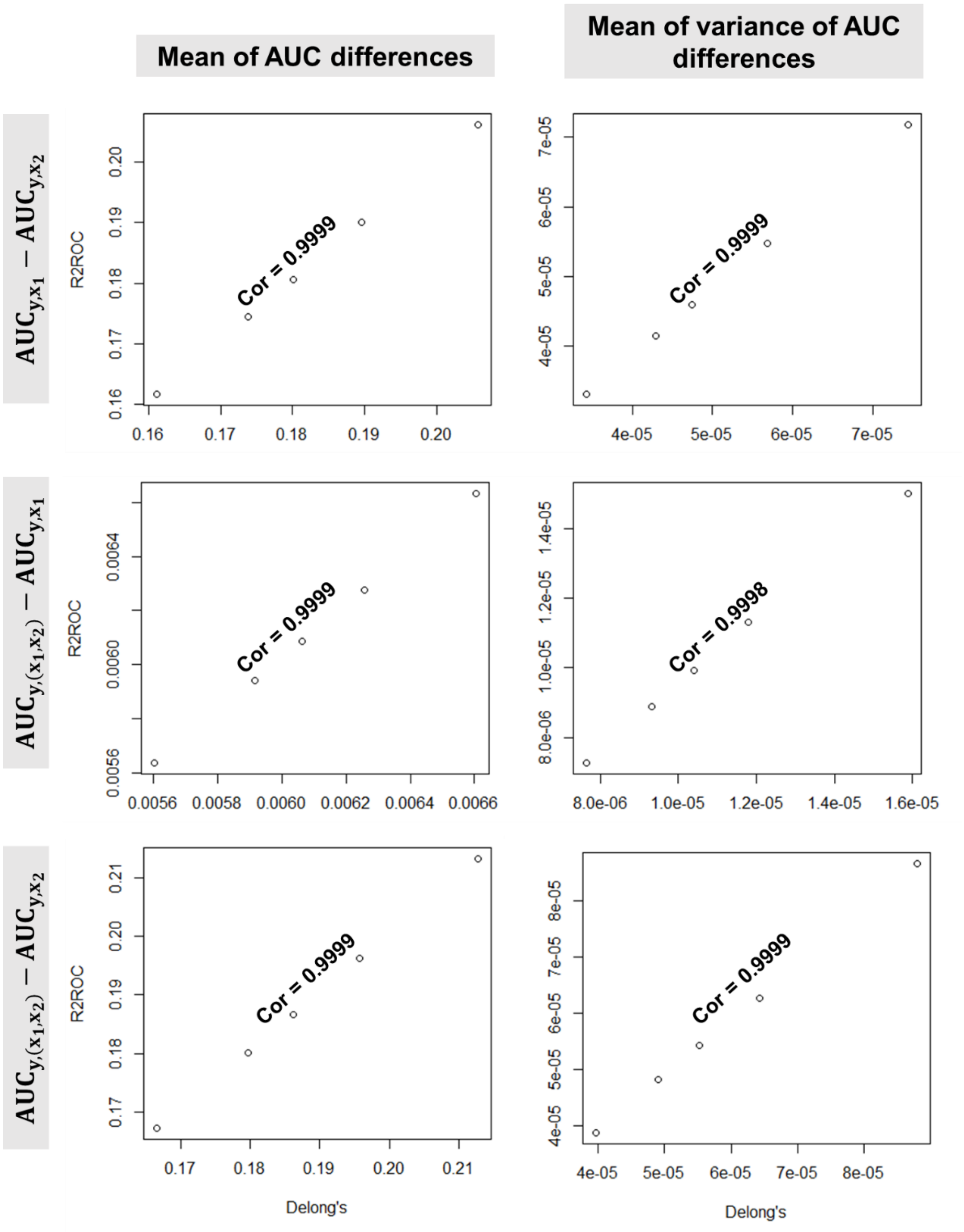


**Figure S3: Agreement between DeLong's and R2ROC methods for AUC difference and estimated variance of AUC difference when varying population prevalence levels.**

The dots represent the averages of AUC differences and estimated variances of AUC difference calculated from 10,000 simulated replicates, revealing the strong agreement between DeLong's and R2ROC methods across various population prevalence levels (*k*=0.05, 0.10, 0.15, 0.20, and 0.50). The comparison involves both non-nested ($AUC_{y,x_1} - AUC_{y,x_2}$) and nested PRS models ($AUC_{y,(x_1,x_2)} - AUC_{y,x_1}$ and $AUC_{y,(x_1,x_2)} - AUC_{y,x_2}$). To generate the dependent variable (y) representing liability values, as well as the PRSs ($x_1$ and $x_2$), a multivariate normal distribution was employed. The correlation structure was determined by the matrix $\begin{bmatrix} 1 & r_{y,x_1} & r_{y,x_2} \\ r_{y,x_1} & 1 & r_{x_1,x_2} \\ r_{y,x_2} & r_{x_1,x_2} & 1 \end{bmatrix} = \begin{bmatrix} 1 & 0.45 & 0.10 \\ 0.45 & 1 & 0.43 \\ 0.10 & 0.43 & 1 \end{bmatrix}$, mimicking the real correlation structure between UKBB and BBJ PRSs for diabetes. Each simulation involved 30,000 individuals to ensure robustness and statistical power.

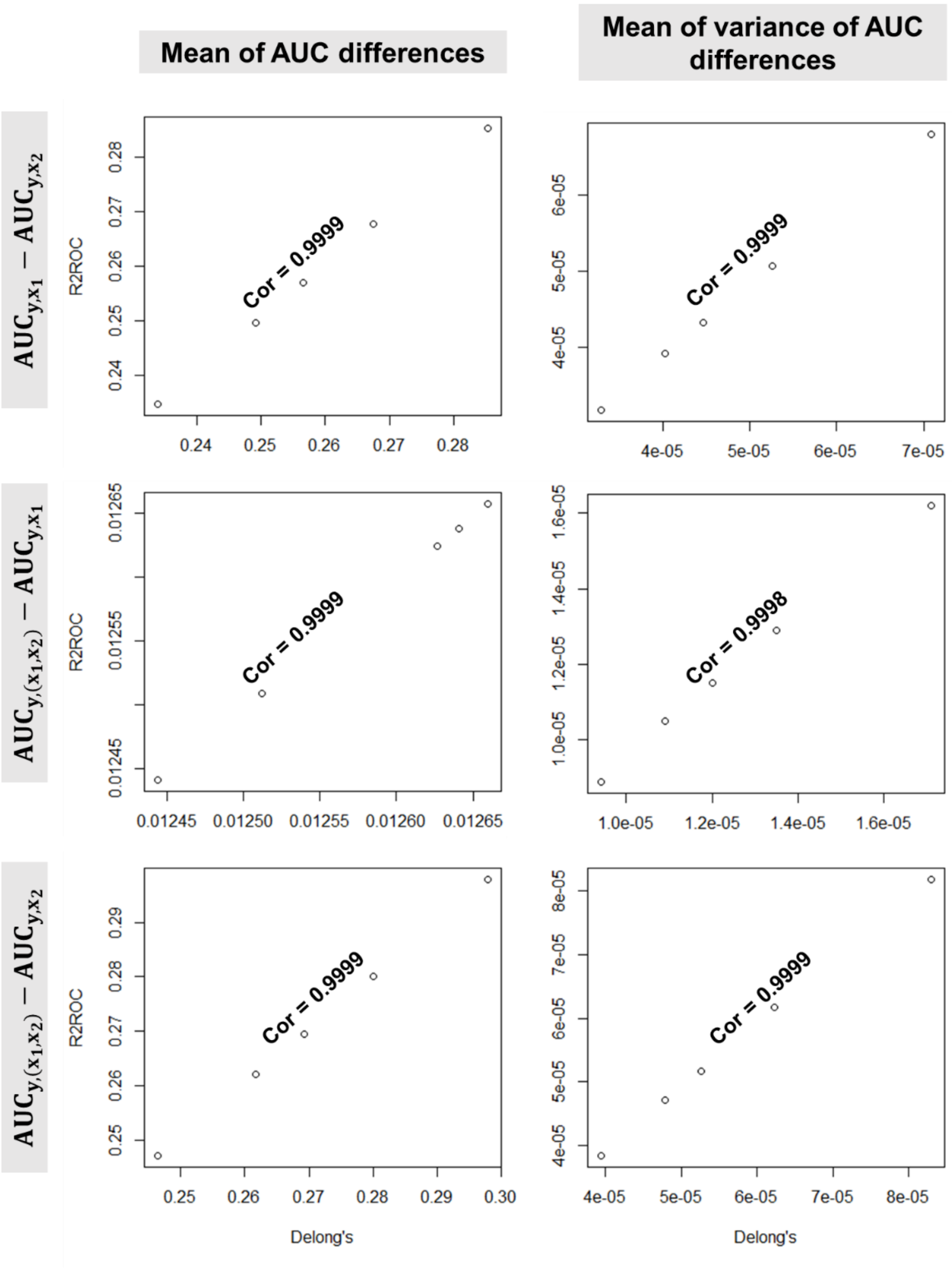


**Figure S4: Agreement between DeLong's and R2ROC methods for AUC difference and estimated variance of AUC difference when varying population prevalence levels.**

The dots represent the averages of AUC differences and estimated variances of AUC difference calculated from 10,000 simulated replicates, revealing the strong agreement between DeLong's and R2ROC methods across various population prevalence levels (*k*=0.05, 0.10, 0.15, 0.20, and 0.50). The comparison involves both non-nested ($AUC_{y,x_1} - AUC_{y,x_2}$) and nested PRS models ($AUC_{y,(x_1,x_2)} - AUC_{y,x_1}$ and $AUC_{y,(x_1,x_2)} - AUC_{y,x_2}$). To generate the dependent variable (y) representing liability values, as well as the PRSs ($x_1$ and $x_2$), a multivariate normal distribution was employed. The correlation structure was determined by the matrix $\begin{bmatrix} 1 & r_{y,x_1} & r_{y,x_2} \\ r_{y,x_1} & 1 & r_{x_1,x_2} \\ r_{y,x_2} & r_{x_1,x_2} & 1 \end{bmatrix} = \begin{bmatrix} 1 & 0.60 & 0.10 \\ 0.60 & 1 & 0.43 \\ 0.10 & 0.43 & 1 \end{bmatrix}$, mimicking the real correlation structure between UKBB and BBJ PRSs for diabetes. Each simulation involved 30,000 individuals to ensure robustness and statistical power.

### Supplemental tables

**Table S1: The comparison between DeLong's and R2ROC methods for AUC difference, estimated variance and observed difference of AUC difference when comparing non-nested PRS models ($\mathrm{AUC}_{y,x_1} - \mathrm{AUC}_{y,x_2}$).**

| Correlation ($r_{y,x_1}$) | Prevalence | DeLong's Method | | | Proposed method (R2ROC) | | |
|---|---|---|---|---|---|---|---|
| | | AUC diff | Variance of diff. | Observed var of diff. | AUC diff | Variance of diff. | Observed var of diff |
| **$r_{y,x_1}$=0.15** | **0.05** | 0.030414 | 6.64E-05 | 6.69E-05 | 0.030425 | 6.20E-05 | 6.24E-05 |
| | **0.10** | 0.027442 | 3.56E-05 | 3.52E-05 | 0.027486 | 3.32E-05 | 3.27E-05 |
| | **0.15** | 0.025765 | 2.52E-05 | 2.55E-05 | 0.025775 | 2.35E-05 | 2.38E-05 |
| | **0.20** | 0.024725 | 2.01E-05 | 2.00E-05 | 0.024757 | 1.88E-05 | 1.88E-05 |
| | **0.50** | 0.02258 | 1.29E-05 | 1.29E-05 | 0.0226 | 1.21E-05 | 1.19E-05 |
| **$r_{y,x_1}$=0.30** | **0.05** | 0.11996 | 6.23E-05 | 6.22E-05 | 0.12012 | 5.95E-05 | 5.75E-05 |
| | **0.10** | 0.10915 | 3.38E-05 | 3.36E-05 | 0.10934 | 3.23E-05 | 3.16E-05 |
| | **0.15** | 0.10308 | 2.41E-05 | 2.45E-05 | 0.10328 | 2.30E-05 | 2.30E-05 |
| | **0.20** | 0.09906 | 1.94E-05 | 1.93E-05 | 0.09926 | 1.84E-05 | 1.81E-05 |
| | **0.50** | 0.09099 | 1.26E-05 | 1.27E-05 | 0.09123 | 1.19E-05 | 1.18E-05 |
| **$r_{y,x_1}$=0.45** | **0.05** | 0.2058 | 5.60E-05 | 5.53E-05 | 0.2062 | 5.56E-05 | 5.15E-05 |
| | **0.10** | 0.1896 | 3.08E-05 | 3.22E-05 | 0.19 | 3.06E-05 | 3.00E-05 |
| | **0.15** | 0.1801 | 2.22E-05 | 2.24E-05 | 0.1806 | 2.20E-05 | 2.11E-05 |
| | **0.20** | 0.1738 | 1.79E-05 | 1.84E-05 | 0.1744 | 1.76E-05 | 1.72E-05 |
| | **0.50** | 0.1611 | 1.18E-05 | 1.17E-05 | 0.1617 | 1.14E-05 | 1.09E-05 |
| **$r_{y,x_1}$=0.60** | **0.05** | 0.2854 | 4.96E-05 | 5.03E-05 | 0.2854 | 5.07E-05 | 4.62E-05 |
| | **0.10** | 0.2675 | 2.73E-05 | 2.77E-05 | 0.2678 | 2.83E-05 | 2.57E-05 |
| | **0.15** | 0.2566 | 1.98E-05 | 1.99E-05 | 0.257 | 2.04E-05 | 1.87E-05 |
| | **0.20** | 0.2491 | 1.60E-05 | 1.62E-05 | 0.2496 | 1.65E-05 | 1.53E-05 |
| | **0.50** | 0.2339 | 1.07E-05 | 1.08E-05 | 0.2347 | 1.07E-05 | 1.01E-05 |

The mean AUC differences and estimated variances calculated from 10,000 simulated replicates for DeLong's and R2ROC methods across various population prevalence levels ($k$=0.05, 0.10, 0.15, 0.20, and 0.50). Note that that observed variance of difference were also calculated from 10,000 simulated replicates. The comparison involves both non-nested ($AUC_{y,x_1} - AUC_{y,x_2}$) PRS models. To generate the dependent variable (y) representing liability values, as well as the PRSs ($x_1$ and $x_2$), a multivariate normal distribution was employed. The correlation structure was determined by the matrix $\begin{bmatrix} 1 & r_{y,x_1} & r_{y,x_2} \\ r_{y,x_1} & 1 & r_{x_1,x_2} \\ r_{y,x_2} & r_{x_1,x_2} & 1 \end{bmatrix} = \begin{bmatrix} 1 & \text{various} & 0.10 \\ \text{various} & 1 & 0.43 \\ 0.10 & 0.43 & 1 \end{bmatrix}$, mimicking the real correlation structure between UKBB and BBJ PRSs for diabetes.

**Table S2: The comparison between DeLong's and R2ROC methods for AUC difference, estimated variance and observed difference of AUC difference when comparing nested PRS models ($\mathbf{AUC_{y,(x_1,x_2)} - AUC_{y,x_1}}$)**

| Correlation ($r_{y,x_1}$) | Prevalence | DeLong's Method | | | Proposed method (R2ROC) | | |
|---|---|---|---|---|---|---|---|
| | | AUC diff | Variance of diff. | Observed var of diff. | AUC diff | Variance of diff. | Observed var of diff |
| $r_{y,x_1}$=0.15 | **0.05** | 0.0033717 | 4.47E-06 | 4.20E-06 | 0.00338 | 3.91E-06 | 3.67E-06 |
| | **0.10** | 0.002973 | 2.31E-06 | 2.20E-06 | 0.002971 | 2.03E-06 | 1.94E-06 |
| | **0.15** | 0.0027514 | 1.62E-06 | 1.62E-06 | 0.0027578 | 1.42E-06 | 1.42E-06 |
| | **0.20** | 0.002603 | 1.28E-06 | 1.23E-06 | 0.002608 | 1.12E-06 | 1.08E-06 |
| | **0.50** | 0.002376 | 8.18E-07 | 8.12E-07 | 0.0023758 | 7.16E-07 | 7.03E-07 |
| $r_{y,x_1}$=0.30 | **0.05** | 0.001147 | 7.55E-07 | 6.95E-07 | 0.0011636 | 6.62E-07 | 6.24E-07 |
| | **0.10** | 0.0010083 | 3.91E-07 | 3.72E-07 | 0.001015 | 3.45E-07 | 3.26E-07 |
| | **0.15** | 0.0009509 | 2.79E-07 | 2.74E-07 | 0.0009572 | 2.46E-07 | 2.43E-07 |
| | **0.20** | 0.0009036 | 2.22E-07 | 2.15E-07 | 0.0009114 | 1.96E-07 | 1.93E-07 |
| | **0.50** | 0.0008264 | 1.43E-07 | 1.41E-07 | 0.00083 | 1.26E-07 | 1.22E-07 |
| $r_{y,x_1}$=0.45 | **0.05** | 0.006606 | 2.54E-06 | 2.53E-06 | 0.006634 | 2.26E-06 | 2.26E-06 |
| | **0.10** | 0.006257 | 1.44E-06 | 1.40E-06 | 0.006275 | 1.30E-06 | 1.28E-06 |
| | **0.15** | 0.006063 | 1.07E-06 | 1.09E-06 | 0.006087 | 9.64E-07 | 9.87E-07 |
| | **0.20** | 0.005917 | 8.74E-07 | 8.69E-07 | 0.005942 | 7.90E-07 | 7.88E-07 |
| | **0.50** | 0.005604 | 5.90E-07 | 5.86E-07 | 0.005636 | 5.30E-07 | 5.30E-07 |
| $r_{y,x_1}$=0.60 | **0.05** | 0.012444 | 2.94E-06 | 2.92E-06 | 0.012295 | 2.66E-06 | 2.62E-06 |
| | **0.10** | 0.012641 | 1.84E-06 | 1.83E-06 | 0.012477 | 1.70E-06 | 1.67E-06 |
| | **0.15** | 0.01266 | 1.43E-06 | 1.44E-06 | 0.012509 | 1.32E-06 | 1.32E-06 |
| | **0.20** | 0.012627 | 1.21E-06 | 1.19E-06 | 0.012498 | 1.11E-06 | 1.10E-06 |
| | **0.05** | 0.012512 | 8.68E-07 | 8.87E-07 | 0.012409 | 7.80E-07 | 7.93E-07 |

The mean AUC differences and estimated variances calculated from 10,000 simulated replicates for DeLong's and R2ROC methods across various population prevalence levels ($k$=0.05, 0.10, 0.15, 0.20, and 0.50). Note that that observed variance of difference were also calculated from 10,000 simulated replicates. The comparison involves both nested ($AUC_{y,(x_1,x_2)} - AUC_{y,x_1}$) PRS models. To generate the dependent variable (y) representing liability values, as well as the PRSs ($x_1$ and $x_2$), a multivariate normal distribution was employed. The correlation structure was determined by the matrix $\begin{bmatrix} 1 & r_{y,x_1} & r_{y,x_2} \\ r_{y,x_1} & 1 & r_{x_1,x_2} \\ r_{y,x_2} & r_{x_1,x_2} & 1 \end{bmatrix} = \begin{bmatrix} 1 & \text{various} & 0.10 \\ \text{various} & 1 & 0.43 \\ 0.10 & 0.43 & 1 \end{bmatrix}$, mimicking the real correlation structure between UKBB and BBJ PRSs for diabetes.

**Table S3: The comparison between DeLong's and R2ROC methods for AUC difference, estimated variance and observed difference of AUC difference when comparing nested PRS models ($\mathbf{AUC_{y,(x_1,x_2)} - AUC_{y,x_2}}$)**

| Correlation ($r_{y,x_1}$) | Prevalence | DeLong's Method | | | Proposed method (R2ROC) | | |
|---|---|---|---|---|---|---|---|
| | | AUC diff. | Variance of diff. | Observed var of diff. | AUC diff. | Variance of diff. | Observed var of diff. |
| $r_{y,x_1}$=0.15 | **0.05** | 0.03375 | 4.27E-05 | 4.21E-05 | 0.03375 | 3.91E-05 | 3.86E-05 |
| | **0.10** | 0.03039 | 2.27E-05 | 2.28E-05 | 0.03039 | 2.07E-05 | 2.08E-05 |
| | **0.15** | 0.02854 | 1.61E-05 | 1.62E-05 | 0.02854 | 1.47E-05 | 1.48E-05 |
| | **0.20** | 0.02733 | 1.28E-05 | 1.26E-05 | 0.02736 | 1.17E-05 | 1.15E-05 |
| | **0.50** | 0.02497 | 8.24E-06 | 8.18E-06 | 0.02499 | 7.53E-06 | 7.52E-06 |
| $r_{y,x_1}$=0.30 | **0.05** | 0.12148 | 7.31E-05 | 7.46E-05 | 0.12163 | 7.05E-05 | 7.03E-05 |
| | **0.10** | 0.11019 | 3.93E-05 | 3.89E-05 | 0.1104 | 3.79E-05 | 3.69E-05 |
| | **0.15** | 0.10402 | 2.81E-05 | 2.84E-05 | 0.10425 | 2.69E-05 | 2.69E-05 |
| | **0.20** | 0.09993 | 2.25E-05 | 2.26E-05 | 0.10015 | 2.16E-05 | 2.15E-05 |
| | **0.50** | 0.09184 | 1.46E-05 | 1.48E-05 | 0.09208 | 1.39E-05 | 1.39E-05 |
| $r_{y,x_1}$=0.45 | **0.05** | 0.2128 | 7.54E-05 | 7.75E-05 | 0.2132 | 7.62E-05 | 7.50E-05 |
| | **0.10** | 0.1958 | 4.14E-05 | 4.12E-05 | 0.1962 | 4.17E-05 | 3.94E-05 |
| | **0.15** | 0.1862 | 2.99E-05 | 3.06E-05 | 0.1867 | 2.99E-05 | 2.94E-05 |
| | **0.20** | 0.1797 | 2.42E-05 | 2.41E-05 | 0.1802 | 2.40E-05 | 2.33E-05 |
| | **0.50** | 0.1666 | 1.60E-05 | 1.58E-05 | 0.1673 | 1.56E-05 | 1.50E-05 |
| $r_{y,x_1}$=0.60 | **0.05** | 0.298 | 6.86E-05 | 6.89E-05 | 0.2979 | 7.20E-05 | 6.69E-05 |
| | **0.10** | 0.28 | 3.83E-05 | 3.88E-05 | 0.2801 | 4.03E-05 | 3.80E-05 |
| | **0.15** | 0.2692 | 2.80E-05 | 2.76E-05 | 0.2695 | 2.92E-05 | 2.67E-05 |
| | **0.20** | 0.2617 | 2.29E-05 | 2.29E-05 | 0.2621 | 2.36E-05 | 2.22E-05 |
| | **0.05** | 0.2464 | 1.54E-05 | 1.55E-05 | 0.2471 | 1.54E-05 | 1.48E-05 |

The mean AUC differences and estimated variances calculated from 10,000 simulated replicates for DeLong's and R2ROC methods across various population prevalence levels ($k$=0.05, 0.10, 0.15, 0.20, and 0.50). Note that that observed variance of difference were also calculated from 10,000 simulated replicates. The comparison involves both nested ($AUC_{y,(x_1,x_2)} - AUC_{y,x_2}$) PRS models. To generate the dependent variable (y) representing liability values, as well as the PRSs ($x_1$ and $x_2$), a multivariate normal distribution was employed. The correlation structure was determined by the matrix $\begin{bmatrix} 1 & r_{y,x_1} & r_{y,x_2} \\ r_{y,x_1} & 1 & r_{x_1,x_2} \\ r_{y,x_2} & r_{x_1,x_2} & 1 \end{bmatrix} = \begin{bmatrix} 1 & \text{various} & 0.10 \\ \text{various} & 1 & 0.43 \\ 0.10 & 0.43 & 1 \end{bmatrix}$, mimicking the real correlation structure between UKBB and BBJ PRSs for diabetes.
